# Protein profiling of WERI RB1 and etoposide resistant WERI ETOR reveals new insights into topoisomerase inhibitor resistance in retinoblastoma

**DOI:** 10.1101/2022.02.19.479927

**Authors:** Vinodh Kakkassery, Timo Gemoll, Miriam M. Krämer, Thorben Sauer, Aysegül Tura, Mahdy Ranjbar, Salvatore Grisanti, Stephanie C. Joachim, Stefan Mergler, Jacqueline Reinhard

## Abstract

Chemotherapy resistance is one of the reasons for eye loss in patients with retinoblastoma (RB). RB chemotherapy resistance has been studied in different cell culture models such as WERI RB1. In addition, chemotherapy resistant RB subclones like the etoposide resistant WERI ETOR cell line have been established to improve the understanding of chemotherapy resistance in RB. The objective of this study was to characterize cell line models of an etoposide sensitive WERI RB1 and its etoposide resistant subclone WERI ETOR by proteomic analysis. Subsequently, quantitative proteomic data served for correlation analysis with known drug perturbation profiles. Methodically, WERI RB1 and WERI ETOR were cultured and prepared for quantitative mass spectrometry (MS). This was carried out in a data-independent acquisition (DIA) mode (Sequential Window Acquisition of All Theoretical Mass Spectra, SWATH-MS). The raw SWATH files were processed using neural networks in a library free mode along with machine learning algorithms. Pathway enrichment was performed using the REACTOME pathway resource and correlated to the Molecular Signature Database (MSigDB) hallmark gene set collections for functional annotation. Furthermore, a drug connectivity analysis using the L1000 database was used to correlate the mechanism-of-action (MOA) for different anticancer reagents to WERI RB1/WERI ETOR signatures. A total of 4,756 proteins were identified across all samples, showing a distinct clustering between the groups. Of these proteins, 64 were significantly altered (q < 0.05 & log2FC |>2|, 22% higher in WERI ETOR). Pathway analysis revealed an enriched metabolic pathway for “retinoid metabolism and transport” in WERI ETOR and for “sphingolipid *de novo* biosynthesis” in WERI RB1. In addition, this study revealed similar protein signatures of topoisomerase inhibitors in WERI ETOR as well as ATPase inhibitors, acetylcholine receptor antagonists and vascular endothelial growth factor receptor (VEGFR) inhibitors in WERI RB1. In this study, WERI RB1 and WERI ETOR were analyzed as a cell line model for chemotherapy resistance in RB using data-independent MS. The global proteome identified activation of “sphingolipid *de novo* biosynthesis” in WERI RB1 and revealed future potential treatment options for etoposide resistance in RB.

## 1. Introduction

To date, retinoblastoma (RB) has an incidence of 1/15,000 to 1/20,000 live births and around 9,000 new cases are diagnosed worldwide every year (1-4). This makes it the most common malignant pediatric ocular tumor globally. Due to an improvement of therapeutic strategies in recent years, the survival rate in western countries is almost 99%, while death rates in Africa and Asia are still high (5-7). Unfortunately, some affected eyes need to be removed due to chemotherapy failure, also caused by chemotherapy resistance (8). Both types of RB, induced either by the spontaneous inactivation of both RB1 alleles or by the combination of one inherited inactive and a second spontaneously inactivated allele, represent an excellent model for RB1 protein eliminated tumors in general.

Chemotherapy resistance mechanisms in RB are unclear so far. To further investigate the mechanisms in RB, the WERI RB1 cell line (Leibniz Institute-German Collection of Microorganisms and Cell Cultures, DSMZ No. ACC90) has been established by spontaneous outgrowth of an enucleated RB eye as published by McFall et al. (9, 10). Among others, Busch et al. characterized the growth behavior of the WERI RB1 cell line under cell culture conditions (11). To identify the mechanisms of chemotherapy resistance, the etoposide resistant subclone WERI ETOR was established by harvesting surviving cells after incubation with increasing etoposide doses from WERI RB1 as described by Stephan et al. (12). Previous studies also examined the parental WERI RB1 and the etoposide resistant subclone WERI ETOR on the molecular level (13-18). Furthermore, Busch et al. demonstrated a higher proliferation rate in WERI ETOR as well as an increased tumor formation and tumor size in an *in vivo* chick chorioallantoic membrane assay (17). Mergler et al. and Oronowicz et al. showed alterations in thermosensitive transient receptor potential channels in WERI ETOR cells (13, 16). A publication by Kakkassery et al. proved an upregulation of sphingosine-1-phosphate in the WERI ETOR cell line as a potential resistance mechanism in RB (14). Recently, Reinhard et al. detected changes in the extracellular matrix (ECM) of WERI ETOR RB cells (15).

Interestingly, the nature of the global proteome in RB remains unclear. As represented by the central dogma of life, proteins are regulated downstream by genetic expression and characterize biochemical signals and treatments of a given cell (19). As such, proteins are perfectly suited to determine the signature of a physiological phenotype as well as the point of intervention for drug and health treatments. Hereby, LC-ESI-MS/MS in a data-independent (DIA/SWATH) mode plays a decisive role. DIA/SWATH uses variable mass windows to enable the complete measurement of all detectable proteins in a sample. This allows the identification and quantification of thousands of proteins in just one measurement. Therefore, the specific aim of this study was to characterize the differences between the cell lines WERI RB1 and WERI ETOR on the proteomic level. The data for this investigation were collected by quantitative mass spectrometry (MS) in DIA mode in combination with machine-learning algorithms for mass spectrometric and statistical evaluation. The generated quantitative proteome data were used to gain further insight into the functional annotation, including intracellular signaling pathways that differ between WERI RB1 and WERI ETOR. Furthermore, a drug connectivity analysis using the L1000 database was used to correlate the mechanism-of-action (MOA) for different anticancer reagents to WERI RB1/WERI ETOR signatures.

## 2. Results

### 2.1 Protein Expression Profiles of WERI RB1 and WERI ETOR Cell Lines

We performed quantitative MS using a DIA mode to detect proteins differentially expressed in WERI RB1 and WERI ETOR cell lines. Using a neuronal-network based identification workflow (DIA-NN, (20)), 4,756 protein groups were identified. All entries with more than 70% missing values (MV) were removed, resulting in 2.2% MVs. The remaining MVs were imputed via replacement by random draws from a normal distribution.

Phenotypic differences between both cell cultures were compared in a t-distributed stochastic neighbor embedding (tSNE) applying protein expression data. tSNE plot showed a clear distinction of the analyzed groups (Figure 1).

**Figure 1.**
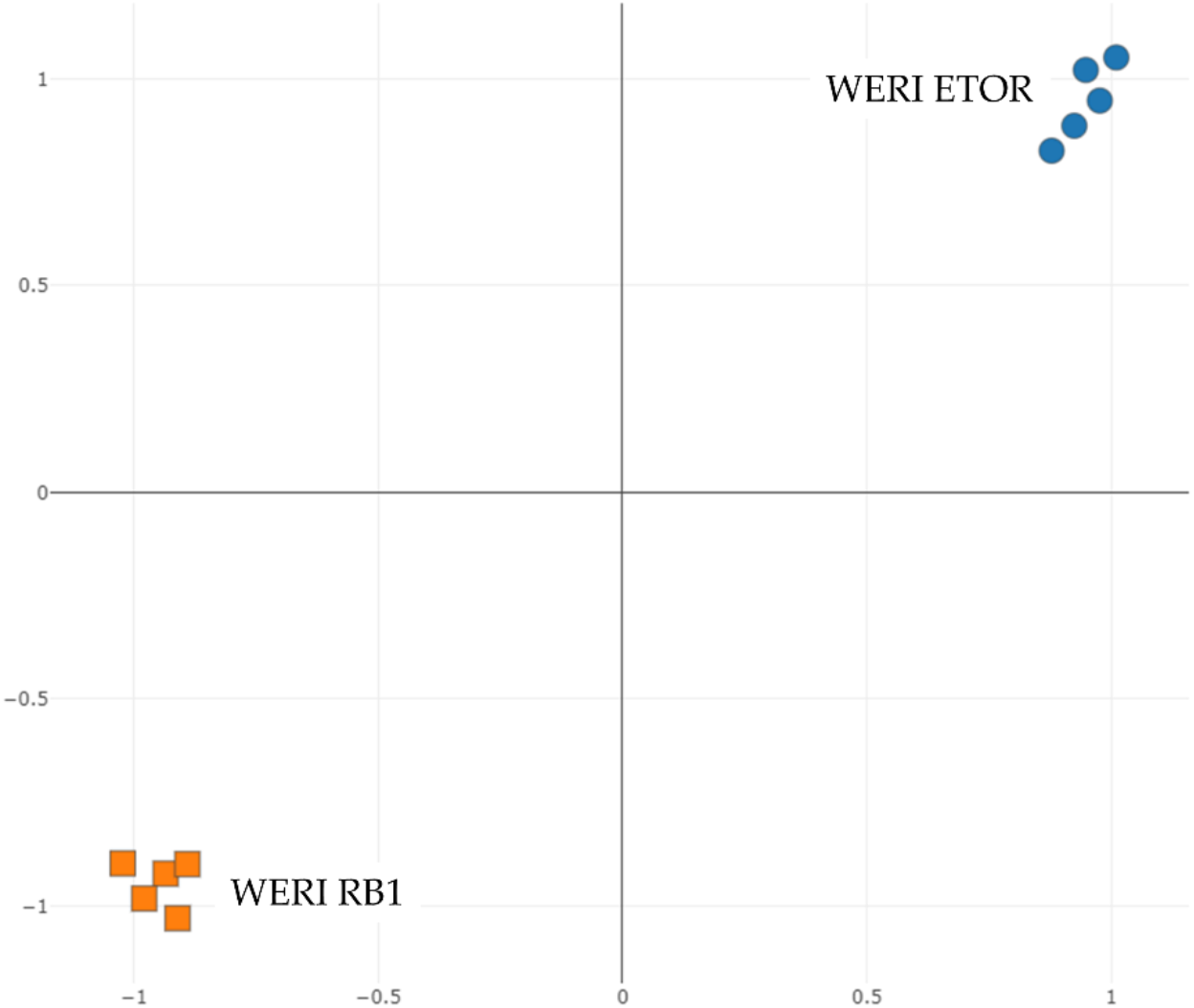
Unsupervised tSNE plot of WERI RB1 (orange) and WERI ETOR (blue) cell line samples. The plot visualizes the close relationship with cell lines and the distinct clustering between sample groups (n=5).

In total, 64 differentially expressed proteins (q < 0.05 & log2FC |>2|) between the two groups were found. Of these proteins, 22 (34%) showed a higher concentration, while 42 (66%) had a lower concentration in the WERI ETOR cell line (Figure 2). The protein with the highest differential expression in WERI ETOR was dehydrogenase/reductase 2 (DHRS2; q-value: 0.0000305 & log2FC: 4.06), whereas H1.5 linker histone, cluster member (H1-5) showed the highest differential expression in WERI RB1 (q-value: 0.00000285 & log2FC: -7.81).

**Figure 2.**
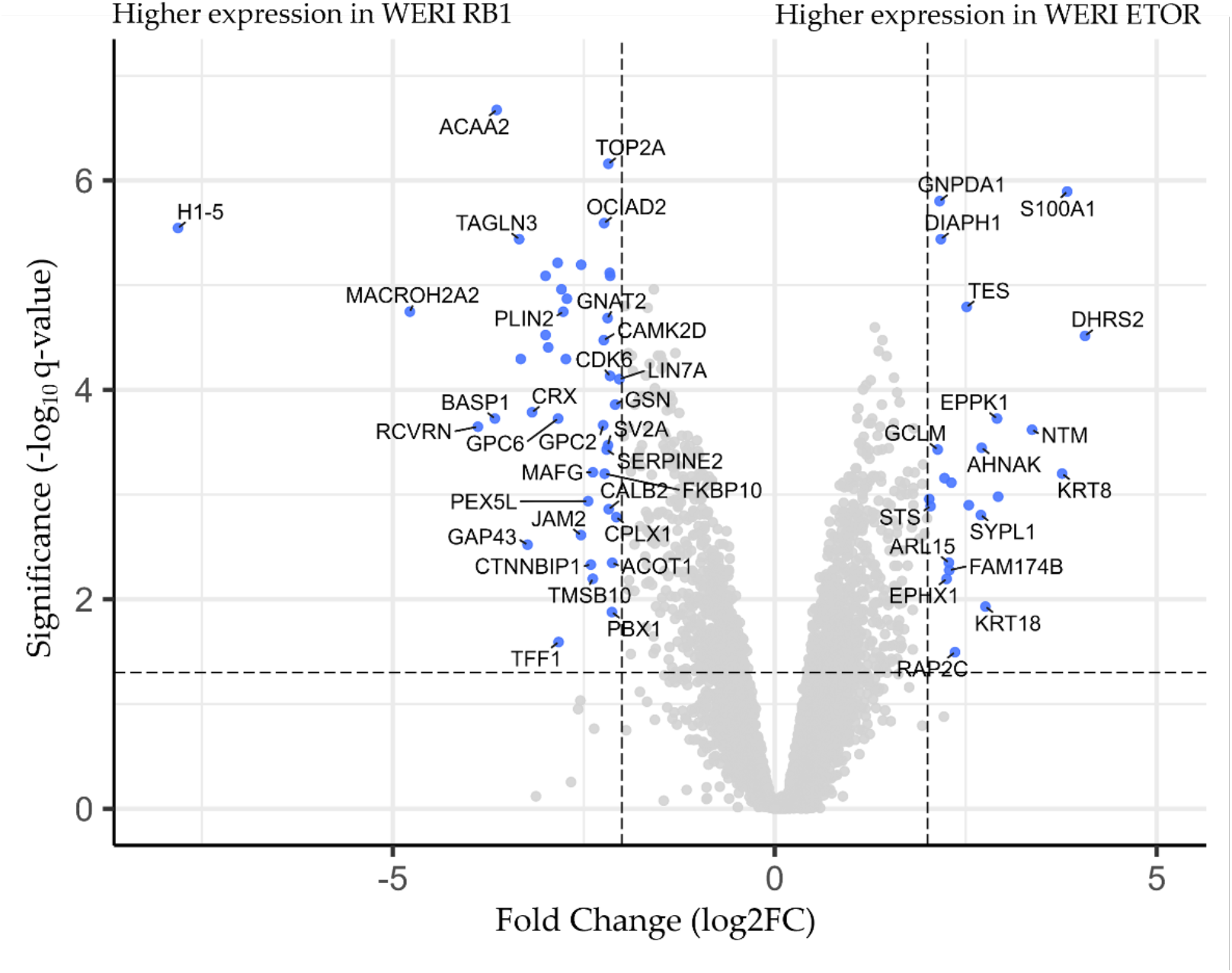
Volcano-plot visualizing fold-change (x-axis) and statistical significance (y-axis). Proteins with a higher concentration in the WERI ETOR cell line are presented on the right side. Proteins with a lower concentration in the WERI ETOR cell line are displayed on the left side. All blue marked proteins demonstrated a logarithmic fold-change (log2FC) of |>2| and a q-value < 0.05.

For validation of differential expressed proteins, a feature importance calculation based on machine learning algorithms (LASSO, elastic nets, random forest & extreme gradient boosting) selected best possible proteins (n = 40) to predict group association (Table 1). What stands out in table is that all top important features showed also statistical significance. Additionally, topmost important proteins are displayed using a heatmap clustering (Figure 3).

**Table 1.** Top 40 differentially expressed proteins between WERI RB1 and WERI ETOR cell lines detected by machine learning algorithms (LASSO, elastic nets, random forests, extreme gradient boosting). Proteins are sorted according to their fold-change. Log fold-change > 0 represent a higher concentration of the protein in the WERI ETOR group. Significance was calculated using four commonly accepted methods in the literature (Welch t-test, limma, edgeR, DESeq2) and merged (meta.q-value).

| Gene Symbol | Full protein name | Chromosome | meta.q-value | Log fold-change [WERI ETOR / WERI RB1] |
| --- | --- | --- | --- | --- |
| DHRS2 | Dehydrogenase/reductase SDR family member 2 | 14 | 3.051e-5 | 4.062 |
| S100A1 | Protein S100-A1 | 1 | 1.275e-6 | 3.828 |
| KRT8 | Keratin 8 | 12 | 6.307e-4 | 3.762 |
| NTM | NAC domain-containing protein 69 | 11 | 2.402e-4 | 3.368 |
| EPPK1 | Epiplakin | 8 | 1.879e-4 | 2.910 |
| AHNAK | AHNAK nucleoprotein | 11 | 3.570e-4 | 2.708 |
| SEPTIN6 | Septin 6 | X | 1.262e-3 | 2.540 |
| TES | Testin | 7 | 1.615e-5 | 2.513 |
| DIAPH1 | Protein diaphanous homolog 1 | 5 | 3.634e-6 | 2.174 |
| GNPDA1 | Glucosamine-6-phosphate isomerase 1 | 5 | 1.576e-6 | 2.158 |
| CRLF3 | Cytokine receptor like factor 3 | 17 | 5.155e-4 | 1.942 |
| MEX3A | RNA-binding protein MEX3A | 1 | 4.712e-5 | -1.871 |
| CDH2 | Cadherin-2 | 18 | 1.581e-5 | -1.955 |
| TOP2B | DNA topoisomerase 2-beta | 3 | 8.151e-6 | -2.155 |
| CDK6 | Cyclin dependent kinase 6 | 7 | 7.362e-5 | -2.156 |
| ATP2B1 | Plasma membrane calcium-transporting ATPase 1 | 12 | 7.630e-6 | -2.160 |
| TOP2A | DNA topoisomerase 2-alpha | 17 | 6.932e-7 | -2.178 |
| GNAT2 | Guanine nucleotide-binding protein G(t) subunit alpha-2 | 1 | 2.064e-5 | -2.189 |
| SERPINE2 | Serpin family E member 2 | 2 | 3.716e-4 | -2.201 |
| OCIAD2 | OCIA domain-containing protein 2 | 4 | 2.556e-6 | -2.234 |
| CAMK2D | Calcium dependent protein kinase II | 4 | 3.350e-5 | -2.238 |
| CTNNBIP1 | Catenin beta interacting protein 1 | 1 | 4.668e-3 | -2.408 |
| LMOD1 | Leiomodin-1 | 1 | 6.385e-6 | -2.534 |
| ACBD7 | Acyl-CoA-binding domain-containing protein 7 | 10 | 1.349e-5 | -2.720 |
| GDAP1L1 | Ganglioside-induced differentiation-associated protein 1-like 1 | 20 | 5.081e-5 | -2.734 |
| PLIN2 | Perilipin-2 | 9 | 1.793e-5 | -2.769 |
| DPYSL3 | Dihydropyrimidinase like 3 | 5 | 1.096e-5 | -2.792 |
| AMPH | D-alanyl-D-alanine-carboxypeptidase/endopeptidase Amph | 7 | 6.121e-6 | -2.842 |
| MAP2 | Microtubule-associated protein 2 | 2 | 3.928e-5 | -2.966 |
| ATP1B1 | Sodium/potassium-transporting ATPase subunit beta-1 | 1 | 2.994e-5 | -3.000 |
| GNGT2 | Guanine nucleotide-binding protein subunit gamma-T2 | 17 | 8.151e-6 | -3.000 |
| CRX | Cone-rod homeobox protein | 19 | 1.636e-4 | -3.177 |
| GAP43 | Neuromodulin | 3 | 3.011e-3 | -3.236 |
| SH3BGRL | SH3 domain-binding glutamic acid-rich-like protein 3 | X | 5.071e-5 | -3.325 |
| TAGLN3 | Transgelin-3 | 3 | 3.634e-6 | -3.346 |
| ACAA2 | 3-ketoacyl-CoA thiolase | 18 | 2.119e-7 | -3.640 |
| BASP1 | Brain acid soluble protein 1 | 5 | 1.879e-4 | -3.664 |
| RCVRN | Recoverin | 17 | 2.248e-4 | -3.886 |
| MACROH2A2 | Core histone macro-H2A.2 | 10 | 1.791e-5 | -4.777 |
| H1-5 | Histone H1.5 | 6 | 2.846e-6 | -7.812 |

**Figure 3.**
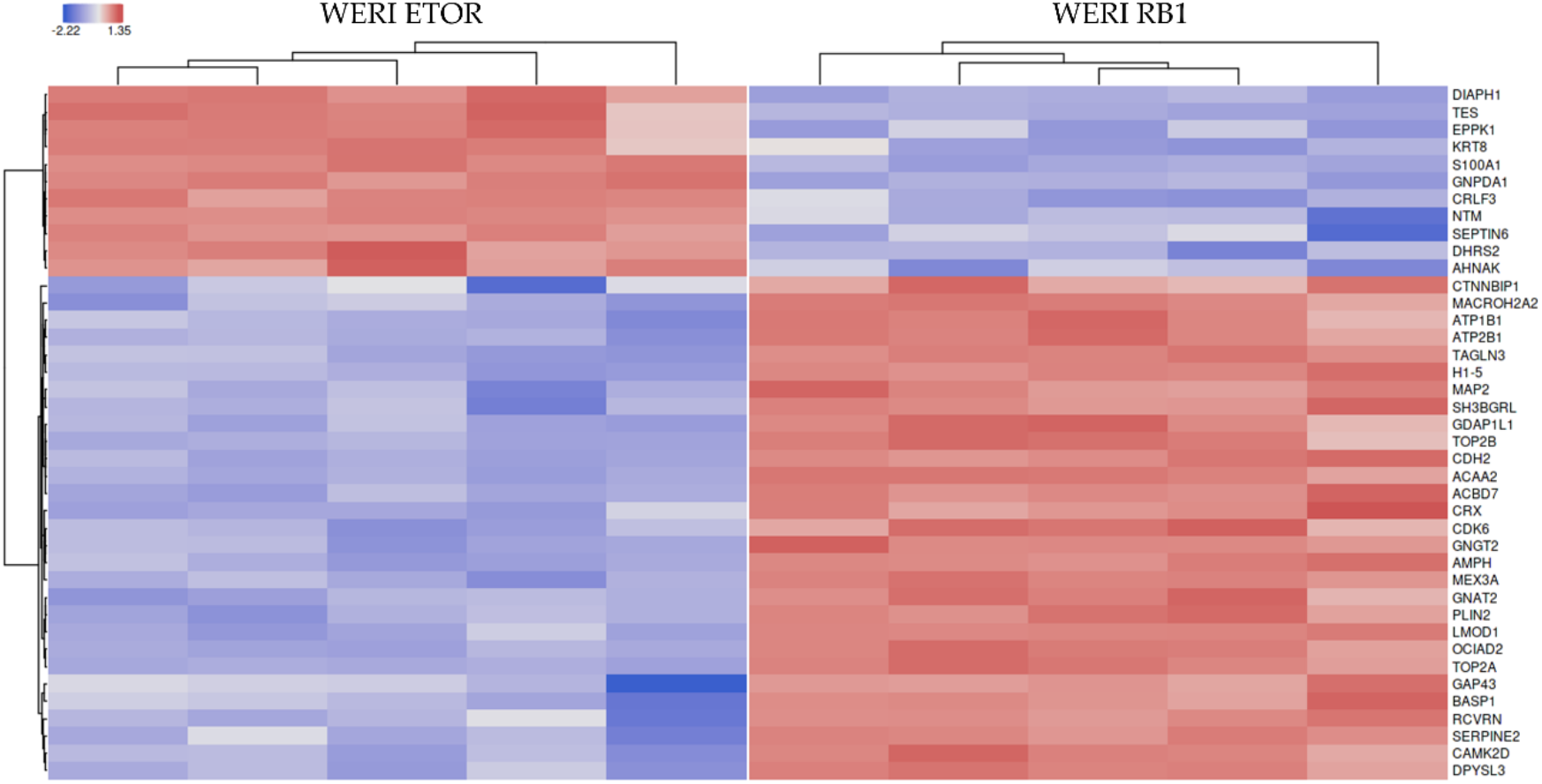
Heatmap of the top 40 selected features according to cumulative ranking by the applied algorithms (LASSO, elastic nets, random forest & extreme gradient boosting). Red colors are high protein expression and blue colors are low protein expression.

### 2.2 Enrichment Pathway Analysis of WERI RB1 and WERI ETOR Protein Signatures

For a deeper insight into the different proteomes of the WERI RB1 and WERI ETOR cell line, all proteins were uploaded to the REACTOME database. REACTOME pathway enrichment algorithm was able to match 3.934 out of 4.756 (83%) proteins which were associated to 2.026 available pathways (submission date: 05-2021). Among others, one of the top enriched metabolic pathways for the WERI ETOR were “retinoid metabolism and transport” (p = 0.0029), while “sphingolipid *de novo* biosynthesis” (p = 0.033) were activated in WERI RB1. Eleven identified proteins were represented in the network of “retinoid metabolism and transport”: Retinol dehydrogenase 11 (RDH11), agrin (AGRN), N(G), N(G)-dimethylarginine dimethylaminohydrolase 1 (DDAH1), apolipoprotein E (APOE), syndecan-1 (SDC1), syndecan-2 (SDC2), glypican-2 (GPC2), apolipoprotein C-III (APOC3), apolipoprotein A-I (APOA1), syndecan-4 (SDC4) and glypican-6 (GPC6). The REACTOME network “sphingolipid *de novo* biosynthesis” contained eight identified proteins, namely aldehyde dehydrogenase family 3 member A2 (ALDH3A2), vesicle-associated membrane protein-associated protein A (VAPA), oxysterol-binding protein 1 (OSBP), serine palmitoyltransferase 1 (SPTLC1), sphingosine-1-phosphate lyase 1 (SGPL1), vesicle-associated membrane protein-associated protein B/C (VAPB), ceramide transfer protein (CERT1) and 3-ketodihydrosphingosine reductase (KDSR). A Voronoi diagram representation of the pathway “metabolism” is shown in Figure 4.

**Figure 4.**
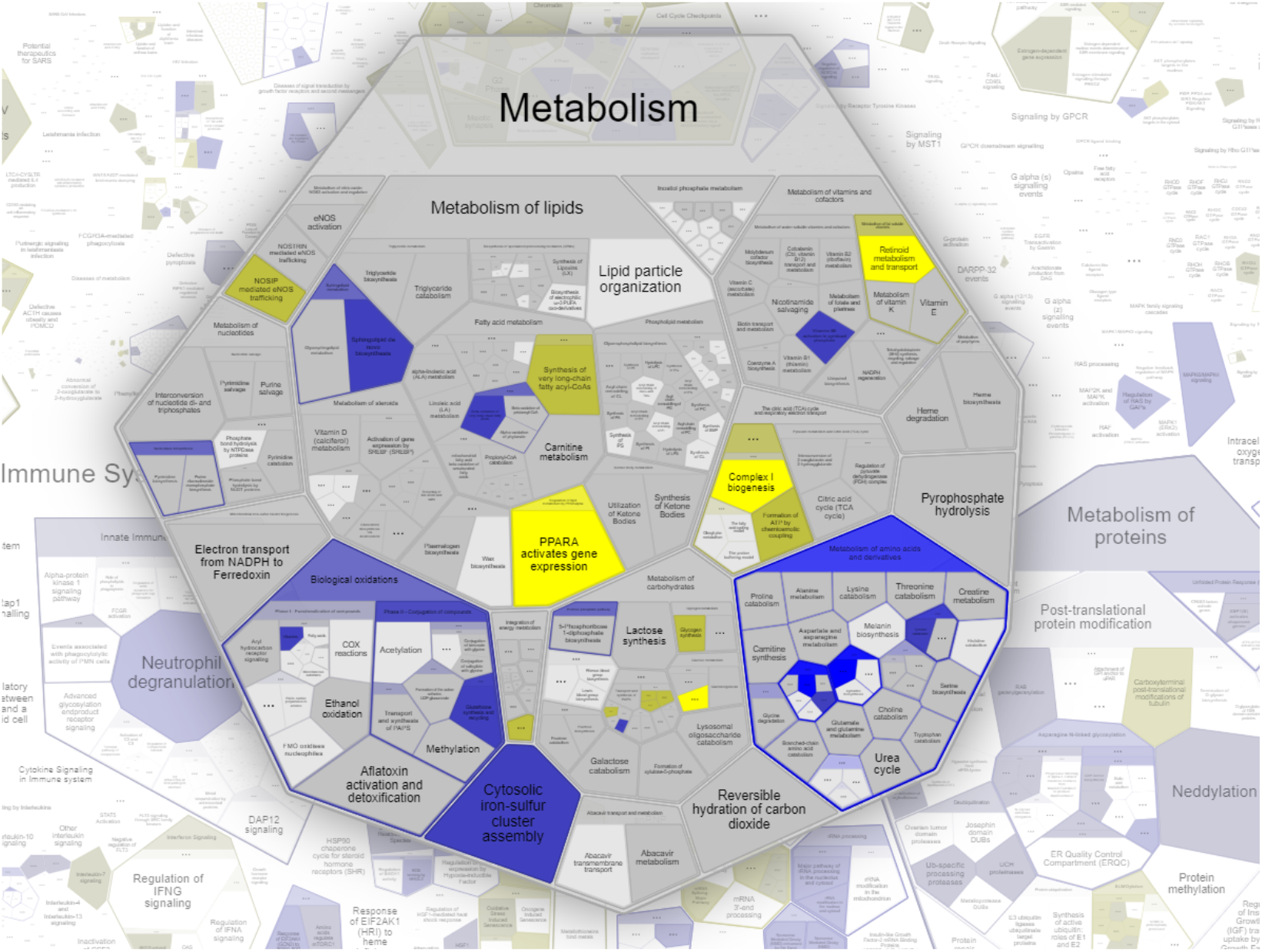
Voronoi diagram representation of the pathway “metabolism” by REACTOME analysis using all expressed protein elements of the WERI RB1 and WERI ETOR cell lines. Yellow colors are pathway activation in WERI RB1; blue colors are pathway activation in WERI ETOR.

### 2.3 Drug Enrichment Analysis of WERI RB1 and WERI ETOR Protein Signatures

To investigate if a certain drug activity or drug sensitivity signature matches with our protein expression data, a drug connectivity analysis using the L1000 database was carried out. The most striking finding of these data was that the correlation between the etoposide resistant cell line WERI ETOR and the parental cell line WERI RB1 revealed a very similar main mechanism of action (MOA) for topoisomerase inhibitors, e.g. etoposide, for the resistant cell line (Figure 5). These data demonstrate that topoisomerase inhibitors (most left) induce specific molecular changes, which are already active in WERI ETOR cells. In turn, topoisomerase inhibitors lose their drug efficacy in these cell lines. ATPase inhibitors, acetylcholine receptor antagonists and VEGFR inhibitors (most right) display the most opposite MOA compared to WERI ETOR cells and should thus overcome the developed therapy resistance.

**Figure 5.**
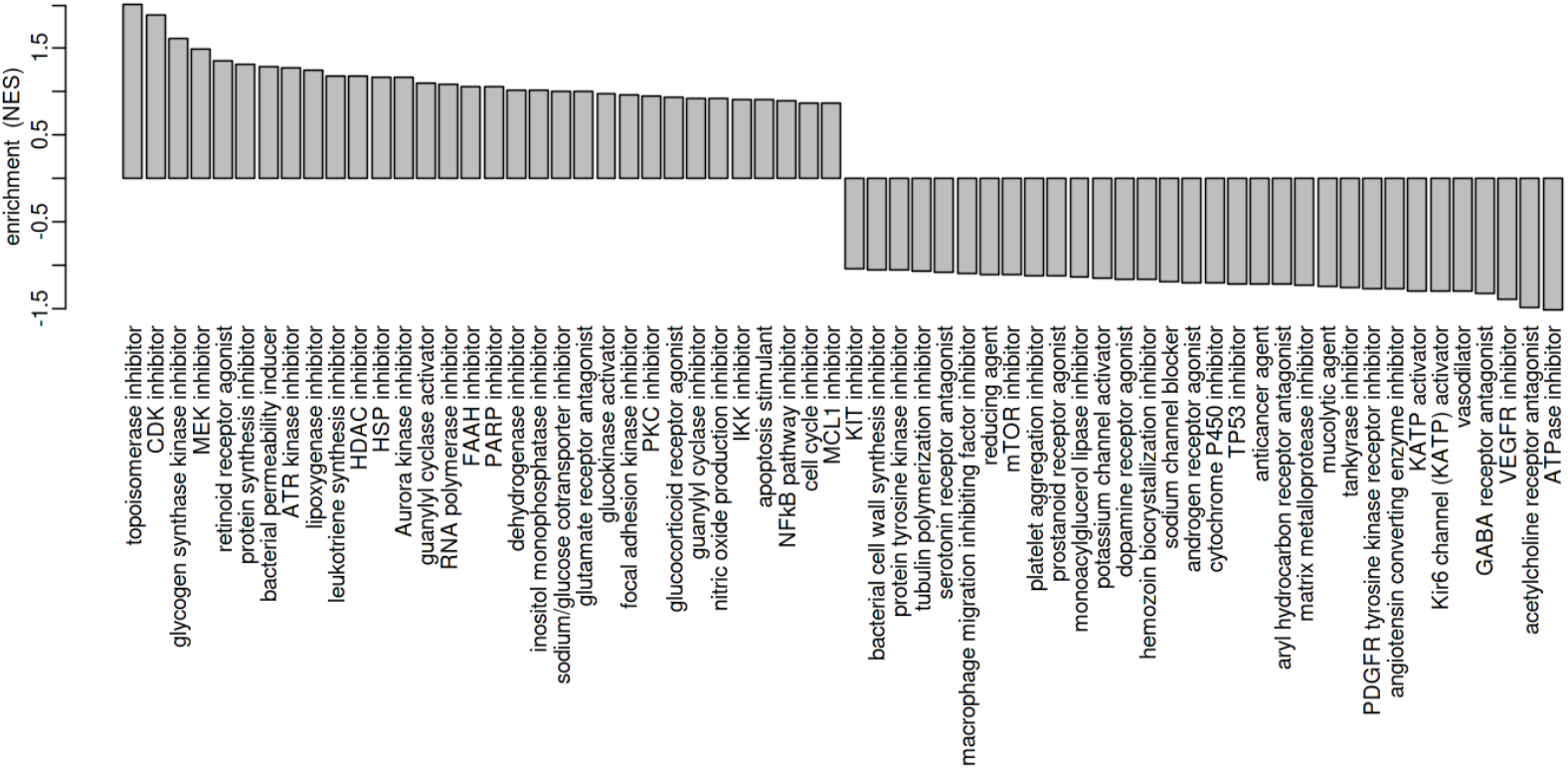
Visualization of mechanism of action (MOA) across enriched drug profiles using the L1000 database. On the vertical axis, GSEA normalized enrichment score of the MOA class is plotted.

## 3. Discussion

Drug resistance in RB is a clinical problem that can lead to enucleation of a child’s eye in order to prevent distant metastasis. The present study was designed to characterize the RB cell line WERI RB1 and its etoposide resistant subclone WERI ETOR at the global proteome level. The comparison of both cell lines identified 64 differentially expressed proteins (log2FC of |>2| and a q-value < 0.05; 22 higher & 42 lower expressed in WERI ETOR). Pathway analysis by REACTOME algorithms revealed the top enriched metabolic pathway “retinoid metabolism and transport” for WERI ETOR, while “sphingolipid *de novo* biosynthesis” was activated in WERI RB1. These results support previous findings that linked RB samples with healthy biospecimens: Danda et al. identified a total of 3,587 proteins using isobaric tags for relative and absolute quantitation (iTRAQ)-based quantitative mass spectrometric analysis and found 899 differentially expressed proteins between RB and healthy human retina samples (21). Using the Ingenuity Pathway Knowledge Database, top identified molecular and cellular processes were lipid metabolism, molecular transportation, small molecule biochemistry, nucleic acid metabolism as well as DNA replication, recombination and repair (21). Cheng et al., on the other hand, analyzed aqueous humour samples from RB patients and compared them with aqueous humour samples from patients without intraocular cancer using the comparative proteomic technique of iTRAQ coupled with offline two-dimensional liquid chromatography-tandem mass spectrometry. A total of 83 proteins were identified to be differentially expressed, of which 44 were upregulated and 39 downregulated in the RB group (22). Assisted by the DAVID bioinformatics resource, protein expression in RB has been implicated in endopeptidase inhibitory activity, peptidase inhibitor, enzyme inhibitor, serine-type endopeptidase inhibitor, structural molecule, lipid binding and carbohydrate binding activities (22). Furthermore, in a different setting, Danda and colleagues performed proteomic analysis on human RB and healthy retina samples. A total of 3,122 proteins were detected in this study, with 282 upregulated and 381 downregulated proteins in the RB sample (23). Using DAVID bioinformatics resource, analysis revealed that most of the regulated proteins were primarily involved in glycoprotein, amyloid acute-inflammatory and defensive responses. Next, Naru et al. analyzed a human papilloma virus positive RB sample, a human papilloma virus negative RB sample and healthy human retina tissue (24). Collectively, 2,828 proteins were identified in this study. While 78 proteins were differentially expressed between the human papilloma virus positive RB sample and the human healthy control retina sample, 88 proteins showed a differentially expression between the human papilloma virus negative RB sample and the human healthy control retina tissue. Bioinformatic analysis performed with the PANTHER classification system showed that most of these proteins were involved in catalytic activity, binding activity, structural molecule, enzyme regulator activity, transporter activity, receptor activity, antioxidant activity, nucleic acid binding transcription factor activity and translation regulator (24). Finally, Galardi et al. performed proteomic profiling by high resolution MS of exosomes from a primary RB tumor cell line and a primary RB cell line established from vitreous seeds, which resulted in the identification of 3,637 proteins (25). Gene enrichment analysis of exclusively and differentially expressed proteins and network analysis detected upregulated proteins related to invasion and metastasis involving extracellular matrix remodeling and interaction, resistance to anoikis and the metabolism of glucose and amino acids in RB vitreous seeds exosomes (25). Taken together, these proteomic analyzes had in common that a more invasive or metastatic RB situation was associated with signaling pathways related to proliferation or inhibition of cell death as well as modulation of the ECM.

Complementary to the previous results, our study identified 4,756 proteins by data-independent MS and revealed differences between the etoposide sensitive WERI RB1 and the etoposide resistant WERI ETOR cell lines. Pathway analysis revealed induction of e.g., “retinoid metabolism and transport” in WERI ETOR cells. This result is consistent with a study of Nicoud et al., which demonstrated that subretinal delivery of Y79 RB cell-infected retinoid-binding protein gene promoters led to formation of photoreceptor cells in the mouse retina (26). In line with this, Vene et al. showed that N-(4-hydroxyphenyl) retinamide inhibited RB tumor growth in a mouse model *in vivo* and induced cell death in Y79 RB cells *in vitro* (27). Khanna et al. detected that retinoic acid upregulates neural retina leucine zipper, a transcription factor expressed in photoreceptors in Y79 (28). In contrast, Kyritsis et al. have demonstrated an inhibitory effect of retinol and retinoid acid on the RB cell line Y79 (29).

Interestingly, as constituents of the “retinoid metabolism and transport” network, we also identified the heparan sulfate proteoglycans (HSPGs) and membrane-linked ECM receptors glypican-2 and -6 as well as syndecan-1, -2 and -4 downregulated in the WERI RB1. As components of the tumor microenvironment, ECM proteins play a critical role in tumor growth and metastasis (30). In this regard, we previously observed an extensive remodeling of various ECM components in the WERI ETOR compared to the WERI RB1 cell line, indicating that the ECM also play a role in mediating chemotherapy resistance formation of RB (15). HSPGs are key players in various processes during neural development and malignant situations (30, 31). They preferentially bind to the basal lamina of blood vessels as well as to ependymal and endothelial surfaces where they act as disposal sites for a variety of cytokines and growth factors (32). Particularly, syndecan-1 and -2 have been described to influence VEGF-dependent neovascularization, an important mechanism to influence RB growth (33-35). Lau et al. observed a reduced glypican-6 mRNA level in RB associated with non-random allelic loss at 13q31 that could contribute to the development of RB (36).

Furthermore, we observed an activation of the “sphingolipid *de novo* biosynthesis” pathway in WERI RB1, which could most likely induce apoptosis and therefore lead to an etoposide vulnerability. Several studies already noted an effect of sphingolipids on cancer (37). A previous study by our group revealed a functional association of the sphingolipid metabolism and WERI ETOR resistance, consistent with the results from our current proteomics study (14). After exposure to etoposide, sphingosine, an apoptotic mediator, was upregulated in WERI RB1 and WERI ETOR, but only in WERI ETOR the anti-apoptotic mediator sphingosine-1-phosphate was upregulated (14). Also, Kim et al. showed that exposure to bisphenol A alters transcriptomic and proteomic dynamics in the RB cell line Y79 (38).

According to these data of pathway activation, we can infer that both “retinoid metabolism and transport” and “sphingolipid *de novo* biosynthesis” may play an important role during therapy resistance in RB. Our results are in line with these studies, which also observed a promotion of retinal cell genesis and may increase resistance of the WERI ETOR.

The most relevant finding in the clinical context was the similar mechanism of action (MOA) of topoisomerase inhibitors but the different MOA of ATPase inhibitors, acetylcholine receptor antagonists and VEGFR inhibitors compared to WERI ETOR. Hence, these findings suggest that biochemical processes of topoisomerase inhibitors as well as WERI ETOR are likely to share the same “positive” molecules including their expression data. It may therefore be the case that etoposide does not affect WERI ETOR cell populations and thus induce no therapeutic activity. Furthermore, and for the same reasons, we speculate that ATPase inhibitors, acetylcholine receptor antagonists and/or VEGFR inhibitors may be more effective treatment options for etoposide resistant RB. To develop a complete picture of new treatment strategies in RB, additional studies will be needed that investigate WERI ETOR and WERI RB1 using drug classes suggested above.

Nevertheless, our study had some limitations. Our *in vitro* cell model only partially mimics chemotherapy resistance in RB *in vivo*. Therefore, to further test our hypothesis regarding the potential mechanism of resistance in RB, experiments in an animal model may be promising as a next step. Also, pathway as well as the MOA enrichment analysis were only examined under untreated conditions for WERI RB1 and WERI ETOR. Along these lines, future experiments should investigate how both cell lines respond to recommended alternative treatment options.

## 4. Materials and Methods

### 4.1 Cell Culture

The etoposide sensitive and resistant RB cell lines WERI RB1 and WERI ETOR were cultivated as previously described (13-17). The identities of these RB cell lines were verified by DNA fingerprinting and profiling using eight different and highly polymorphic short tandem repeat (STR) loci (Leibniz-Institute DSMZ GmbH, Braunschweig, Germany). The generated STR profiles of the cell lines verified a full match of the respective parental reference STR profiles as determined by a search of the cell bank databases ATCC (Manassas, VA, USA), JCRB (Osaka, Japan), RIKEN (Ibaraki, Japan), KCLB (Seoul, Korea) and DSMZ (Braunschweig, Germany). Additionally, the purity of both cell lines was demonstrated by analysis of mitochondrial DNA sequences from mouse, rat as well as Chinese and Syrian hamster cells. Analyzes with a detection limit of 1:10^5^ showed the absence of mitochondrial sequences from foreign species.

### 4.2 Cell Preparation and Mass Spectrometry

The Pierce™ Mass Spec Sample Prep Kit for Cultured Cells (Thermo Scientific, Waltham, MA, USA) was used for the preparation and precipitation of peptides according to the manufacturer’s instructions. 10^6^ cells (n=5/group) were first lysed with a Cell Lysis Buffer and DNA and RNA were enzymatically digested using Nuclease for Cell Lysis. Total protein concentration was subsequently determined using the fluorescence-based EZQ™ Protein Quantitation Kit (Life Technologies, Carlsbad, CA, USA) according to the manufacturer’s protocol. Briefly, protein samples were spotted and fixed onto an assay paper and then stained with the proprietary fluorescence reagent. Fluorescence visualization was done with the Typhoon™ FLA 9000 laser scanner (GE Healthcare, Chicago, IL, USA). Subsequently, 50µg proteins were reduced, alkylated and precipitated. An enzymatic protein digestion was then carried out by adding the Digestion Buffer and Lys-C Protease to the acetone-precipitated protein pellet. The samples were then frozen at -80°C with Trypsin Storage Solution to stop digestion.

The samples were lyophilized, resuspended in solvent A (0.1% FA) to a final concentration of 1 µg/ml and were loaded into the HPLC (Thermo Scientific, Waltham, MA, USA). The reconstituted peptides were first desalted on a trap column (Luna, 5 µm C18 (2), 20 × 0.3 cm; Phenomenex, Torrance, CA, USA) and separated on an analytical column (LC Column, 3 µm C18 (2), 150mm x 0.3mm, Phenomenex, Torrance, CA, USA) using a multi-step gradient of solvent B (0.1% FA in ACN) in solvent A for 90 min at a flow rate of 5µL/min. Mass spectra were acquired on a Triple ToF 5600+ (ScieX, Framingham, MA, USA) in a data-independent (SWATH) acquisition mode. The working parameters of the MS were Ion Spray Voltage Floating at 5000 V; ion source gas 1, 15; ion source gas 2, 0; curtain gas, 30; source temperature, 0°C. The optimized declustering potential (DP) was set at 100 V. The SWATH acquisition parameters were as follow: one 0.49965 sec MS scan (m/z 400–1.250), followed by 100 variable Q1 windows with the size range 4.6-40.9 Da with 1 Da overlap.

### 4.3 SWATH Data and Quantitative Data Processing

The raw SWATH files were processed using the software tool DIA-NN v1.7.16 (Data-Independent Acquisition by Neural Networks) developed by Vadim Demichev et al. (20). The “match between runs” function was used to first develop a spectral library from data-independent acquisition data. The precursor ion generation settings were set to peptide length of 7-52 amino acids, maximum number of missed cleavages to one and maximum number of variable modifications to zero. The precursor and fragment ion m/z range were 350-1.250 m/z.

Precursors that passed the FDR cut-off of 0.01 were grouped to protein-/gene-groups. For those groups consisting of multiple identifiers, the protein groups were reduced to the first listed identifier. The duplicates from a protein-group resembling were removed.

The filtered dataset was further processed in the software tool Perseus (39). MVs were imputed by random numbers drawn from a normal distribution with a width of 0.3 and down shift of 1.8 (default settings) and total matrix imputation mode. The resulting data matrix was used for subsequent statistical analysis.

### 4.4 Data and Statistical Analysis

Protein data were partly analyzed with the software Omics Playground (BigOmics Analytics, v2.8.5) (40). Cluster analyzes were carried out using t-distributed stochastic neighbor embedding (tSNE) algorithm. To increase the statistical reliability of the protein expression differences, we performed the analysis using commonly accepted methods in the literature, including Welch t-test, limma, edgeR as well as DESeq2 and merged the results. Significant differential protein expression was considered at a q-value of < 0.05 and a logarithmic fold-change (log2FC) of |>2|. Additionally, a variable importance score for each protein was calculated using multiple machine-learning algorithms, including LASSO (41), elastic nets (42), random forests (43) and extreme gradient boosting (44). The findings of the machine learning algorithms were used to validate our marker panels derived from classical statistical tests.

### 4.5 Pathway Enrichment and Drug Connectivity Correlation

Pathway enrichment was performed using the REACTOME pathway resource (www.reactome.org, (45)) to understand high-level protein functions and to link information from large molecular data sets. The REACTOME Camera workflow with a focus on metabolic pathways was used to analyze the function of all quantitative data. Protein signatures were further correlated to known drug profiles from the L1000 database (46) by running the gene-set-enrichment-analysis (GSEA) algorithm (47).

## 5. Conclusions

Our study identified over 4,700 proteins in both cell lines examined and further uncovered significant differences between WERI RB1 and WERI ETOR. In particular, the pathways “retinoid metabolism and transport” and “sphingolipid *de novo* biosynthesis” seem to play an important role in etoposide resistance, which could further clarify the understanding of resistance in RB. By drug connectivity analysis using the L1000 database, this study also revealed a similar MOA for topoisomerase inhibitors and WERI ETOR, but a different MOA and potential new treatment options for ATPase inhibitors, acetylcholine receptor antagonists and VEGFR inhibitors. Taken together, our study, which characterized WERI RB1 and WERI ETOR as a cell line model for the human situation in RB, offers new treatment ideas that should be tested in upcoming experiments.

## Author Contributions

Conceptualization, V.K. and J.R.; Formal Analysis, V.K., T.G., T.S. and J.R.; Funding Acquisition, V.K. and M.M.K.; Methodology, V.K., T.G., M.M.K., S.C.J., S.M. and J.R; Writing—Original Draft, V.K., T.G., T.S. and J.R.; Writing—Review & Editing, M.M.K., A.T., M.R., S.G., S.C.J. and S.M. All authors have read and agreed to the published version of the manuscript.

## Acknowledgments

We thank Andreas Faissner for the constant support. We also thank Sabine Kindermann and Sandra Lata for technical assistance.

